# Semaphorin-Plexin signaling influences early ventral telencephalic development and thalamocortical axon guidance

**DOI:** 10.1101/098236

**Authors:** Manuela D. Mitsogiannis, Graham E. Little, Kevin J. Mitchell

## Abstract

**Background:** Sensory processing relies on projections from the thalamus to the neocortex being established during development. Information from different sensory modalities reaching the thalamus is segregated into specialized nuclei, whose neurons then send inputs to cognate cortical areas through topographically defined axonal connections.

Developing thalamocortical axons (TCAs) normally approach the cortex by extending through the subpallium; here, axonal navigation is aided by distributed guidance cues and discrete cell populations, such as the corridor neurons and the internal capsule (IC) guidepost cells. In mice lacking Semaphorin-6A, axons from the dorsal lateral geniculate nucleus (dLGN) bypass the IC and extend aberrantly in the ventral subpallium. The functions normally mediated by Semaphorin-6A in this system remain unknown, but might depend on interactions with Plexin-A2 and Plexin-A4, which have been implicated in other neurodevelopmental processes.

**Methods:** We performed immunohistochemical and neuroanatomical analyses of thalamocortical wiring and subpallial development in *Sema6a* and *Plxna2;Plxna4* null mutant mice and analyzed the expression of these genes in relevant structures.

**Results:** In *Plxna2;Plxna4* double mutants we discovered TCA pathfinding defects that mirrored those observed in *Sema6a* mutants, suggesting that Semaphorin-6A–Plexin-A2/Plexin-A4 signaling might mediate dLGN axon guidance at subpallial level.

In order to understand where and when Semaphorin-6A, Plexin-A2 and Plexin-A4 may be required for proper subpallial TCA guidance, we then characterized their spatiotemporal expression dynamics during early TCA development. We observed that the thalamic neurons whose axons are misrouted in these mutants normally express Semaphorin-6A but not Plexin-A2 or Plexin-A4. By contrast, all three proteins are expressed in corridor cells and other structures in the developing basal ganglia.

This could be consistent with the Plexins acting as guidance signals through Sema6A as a receptor on dLGN axons, and/or with an indirect effect on TCA guidance due to functions in morphogenesis of subpallial intermediate targets. In support of the latter possibility, we observed that in both *Plxna2;Plxna4* and *Sema6a* mutants some IC guidepost cells abnormally localize in correspondence of the ventral path misrouted TCAs elongate into.

**Conclusions:** These findings implicate Semaphorin-6A–Plexin-A2/Plexin-A4 interactions in dLGN axon guidance and in the spatiotemporal organization of guidepost cell populations in the mammalian subpallium.

## Background

The establishment of specific, finely organized neural circuits comprising often distant central nervous system regions is essential for normal brain functioning. Indeed, many neurological and psychiatric disorders have been characterized as potential neurodevelopmental disconnectivity or abnormal connectivity syndromes [1], including autism spectrum disorders [2-7], schizophrenia [8–16], and attention deficit hyperactivity disorder [17–19].

Growing axonal projections in the brain follow spatially complex, yet remarkably stereotyped pathways en route to their final destinations. Forebrain connections thus develop in a stepwise manner: as axons progressively extend, specific growth instructions are provided by guidance factors located at defined ‘decision points’ along axonal paths [20]. The correct spatiotemporal distribution of guidance molecules, supported by the development and proper assembly of intermediate targets, is therefore just as essential as their ultimate effects on growth cones for normal brain wiring [21–23]. While a wealth of knowledge has been gained in the past two decades on axon guidance molecules and their roles in steering developing axons, our understanding of the processes underlying intermediate target formation, guidance cue patterning of axonal pathways, and cue presentation to elongating fibers is still relatively limited [22, 24]. One of these process is the migration at intermediate points of “guidepost cells”, i.e. discrete, specialized cell populations that finely orient growth cones via short-range cues and direct cell–cell contacts. Interestingly, several lines of evidence have pointed to a role of guidance factors also in guidepost cell migration and positioning, and have shown how these molecules can thus affect axonal pathfinding in an indirect manner (reviewed in Squarzoni, Thion [25]).

In the study of the molecular mechanisms involved in the establishment of topographically arranged neural connections, the thalamocortical system constitutes a unique analytical model. Indeed, in the assembly of neural networks between the cortex and the thalamus, axonal sorting is aided by the presence of guidance molecules patterns, cytoarchitecturally-defined permissive pathways, guidepost cell populations at multiple ‘decision points’, and axon-axon interactions.

In the mouse, thalamocortical axons (TCAs) from the dorsal thalamus (dTh) first extend ventrally into the prethalamus, then make a sharp turn at the diencephalic-telencephalic boundary (DTB) and proceed in a dorso-lateral direction through the ventral telencephalon (vTel) to reach the pallial-subpallial boundary (PSPB). As TCAs travel within the subpallium, they then diverge along the rostro-caudal axis according to the position of their final cortical targets [21, 26–28]. Navigation of axons in these early stages, taking place between embryonic day (E) 11-15 [27, 29], has been demonstrated to rely on the presence of intermediate targets and guiding cell populations [21, 27, 30-32], such as the corridor cells and the guidepost cells found in the internal capsule (IC), the axonal bundle containing all reciprocal connections between cortical and subcortical structures.

So far, no specific guidance mechanisms have been identified in axonal pathfinding from each distinct thalamic nucleus. Matching of thalamic axons with appropriate cortical areas appears to emerge from the combinatorial action of several guidance factors and their receptors, expressed in complementary gradients within the dTh and the intermediate targets delineating axonal paths to the neocortex [21, 27]. However, previous studies by our lab have demonstrated a key role of Semaphorin-6A (Sema6A), a member of the semaphorin protein family, in subpallial pathfinding of the visual subset of thalamocortical connections. In *Sema6a* null mice, all dorsal lateral geniculate nucleus (dLGN) axons can be observed to abnormally extend outside the IC and into more ventral areas of the vTel, while projections from other thalamic nuclei develop normally [33–35]. This results in an invasion of the presumptive visual cortex by somatosensory TCAs from the ventrobasal complex (VB) at embryonic stages that persists until early postnatally. A few days after birth, an approximately normal pattern of thalamocortical connectivity is reestablished by dLGN axons navigating to the visual cortex via alternative routes, and outcompeting VB-originated axons.

The exact mechanism by which Sema6A influences TCA guidance in such a specific manner has yet to be elucidated. In the central nervous system, two Sema6A binding partners have been so far identified, Plexin-A2 (PlxnA2) and Plexin-A4 (PlxnA4). Sema6A typically acts as a ligand for PlxnA2 and PlxnA4 in the brain, but it is known to be capable of both forward and reverse signaling with the two Plexin protein family members [36, 37]. Moreover, the effects of Sema6A–PlexinAs binding are highly context-dependent, and have been shown to be modulated by association between these proteins *in cis* [38–41]. Experimental evidence indicates that interactions between Sema6A and PlxnA2/PlxnA4 control several fundamental processes in the establishment of neural circuits, such as axon guidance [42, 43], axonal growth [44], laminar connectivity formation [41, 45], neural cell migration [46, 47], and dendritogenesis [48].

Considering all these findings, both Plexin family members might also be hypothesized to mediate Sema6A–induced responses in early visual thalamic axon guidance. Therefore, in this study we investigated the potential role of Sema6A–PlxnA2/PlxnA4 signaling in subpallial TCA pathfinding by analyzing the phenotype of single and double null mutant mouse lines for the two Plexin genes. In *Plxna2;Plxna4* double mutants we observed a TCA phenotype almost identical to that seen in *Sema6a* mutants. Expression analyses indicate a non-canonical mode of Semaphorin–Plexin interaction-mediated guidance in this case, as both Plexins are not expressed by the misrouted axons, but all proteins are present in the developing subpallium. This suggests either Plexin–Semaphorin reverse signaling taking place, or an indirect effect of Sema6A, PlxnA2 and PlxnA4 on TCA guidance due to earlier functions in ventral forebrain development. A possible indirect guidance role is supported by the finding that a subset of IC guidepost cells is mislocated in both *Sema6a* and *Plxna2;Plxna4* mutants, suggesting that interactions between these molecules are involved in early morphogenesis of subpallial domains delineating TCA pathways.

## Material and methods

### Animals

All animal procedures were performed in accordance with Irish regulations on the use of animals for scientific purposes (Statutory Instrument No. 566 of 2002 and No. 543 of 2012) and institutional guidelines.

All experiments were performed on embryonic brains taken from C57BL/6J mice (wild-type) (Jackson Laboratories), a C57BL/6J strain carrying a null mutation for the *Sema6a* gene, and a *Plxna2;Plxna4* double null mutant C57BL/6J strain. The *Sema6a* mutation was obtained through the insertion of a gene-trap vector in the 17th intron of *Sema6a,* which results in the production of an intracellularly sequestered N-terminal Sema6A portion–β-galactosidase fusion protein (for further details see Leighton et al. [33] and Mitchell et al. [49]). *Plxna2;Plxna4* knockout mice were generated by crossing single mutant lines obtained using gene-targeting strategies described by Suto and colleagues [41] (*Plxna2*), and Yaron and colleagues [50] (*Plxna4*). The *Plxna4* line was originally generated in a CD1 background and backcrossed to C57BL/6J for 10 generations. Mice were maintained and bred in a 12:12 hour light-dark cycle, in a specific pathogen free animal unit.

Pregnant animals for embryo collection were obtained through timed matings. Embryonic age was calculated considering the day of vaginal plug detection as E0.5. For postnatal mice, the day of birth was designated as P0. Mouse brains were dissected in cold PBS and fixed in 4% paraformaldehyde (PFA)/PBS for 2-4 hours at 4° C. Mouse tail samples (2-5 mm) were also collected for DNA extraction and genotyping.

### Genotyping

Tissue samples were digested overnight at 56° C in a 1:100 solution of Proteinase K (Roche) in Boston Buffer (50 mM Tris-HCl pH 8, 2.5 mM EDTA, 50 mM KCl, 0.45% NP-40, and 0.45% Tween 20). The genotype was determined by PCR, using a small aliquot of the digestion solution, a PCR mix (KAPA2G Fast HotStart ReadyMix with dye, KAPA Biosystems) containing dNTPs, a Taq polymerase and a loading dye, and primers for the gene of interest (see Additional file 1 - Table S1).

### *In situ* hybridization

Digoxigenin (DIG)-labeled antisense (as) riboprobes for *in situ* hybridization were produced from plasmid templates linearized overnight at 37° C with appropriate restriction enzymes (see Additional file 1 - Table S2). A transcription reaction mix was prepared using 1 μg of linearized plasmid template, 2 μL of 10x RNA transcription buffer, 2 μL of 10x DIG labeling mix (Roche), 0.5 μL of RNase OUT (40 U/μL, Invitrogen), 20 U of T3 RNA polymerase, and brought to 20 μL volume with dH_2_O. The reaction mix was incubated for 2 hours at 37° C; 1 U of RNase-free DNase I for 15 minutes at 37°C were then added to remove the template, followed by 2 μL of 0.2 M EDTA at pH 8.0 to stop this reaction. Synthesized RNA was precipitated by adding 2.5 μL of 4 M LiCl and 75 μL of ethanol and incubating at −20° C for over an hour, and afterwards pelleted by centrifugation at 4° C (17,000 g or over, 20 minutes). RNA pellets were washed with 150 μL of 70% ethanol via centrifugation at 4° C (17,000 g or over, 2 minutes), briefly let to air-dry, and re-suspended in 1 mM sodium citrate at pH 6.35.

Serial 60 μm thick free-floating brain sections were obtained by slicing brains embedded in 4% w/v RNase free, low melting point agarose/RNase-free phosphate buffered saline (PBS) (obtained by treatment with diethylpyrocarbonate (DEPC)) with a vibratome (VT1000s, Leica), and were collected in DEPC-treated PBS. Sections were washed twice for 5 minutes in DEPC-treated PBS containing 0.1% Tween-20 (PBT). They were then permeabilized by incubation for 30 minutes in 0.5% Triton X-100 and 0.2% Tween-20 in DEPC-treated PBS at room temperature (RT), and additionally washed for 5 minutes in PBT. Next, half of the PBT volume was removed and replaced by pre-hybridization solution (50% deionized formamide, 5x standard saline citrate (SSC), 1% sodium dodecyl sulfate (SDS), 2.5 mg of yeast tRNA, 2.5 mg of heparin in DEPC-treated dH_2_O) pre-warmed at 65° C. After a 10 minutes incubation, sections were further washed with warm pre-hybridization solution for 10 minutes, and then incubated in fresh pre-warmed pre-hybridization solution for at least 1 hour at 65° C. Warm pre-hybridization solution containing 1 g/mL of the RNA probe of choice was next added to each section series for an overnight incubation at 65° C in a humidity chamber (pre-equilibrated with 50% formamide/5x SSC in DEPC-treated dH_2_O).

Following hybridization, three 30 minutes washes in solution I (50% formamide, 5X SSC, 1% SDS) and three 30 minutes washes in solution III (50% formamide, 2X SSC, 0.1% Tween-20) were performed at 65° C. Sections were then washed three times for 5 minutes in Tris-buffered saline (TBS) containing 1% Tween-20 (TBST) at RT, and incubated in blocking buffer (TBST containing 10% heat-inactivated sheep serum) for at least 1 hour. The buffer was subsequently replaced with an alkaline phosphatase (AP) conjugated anti-DIG antibody solution (1:2000 anti-DIG-AP Fab fragment (Roche) in blocking buffer) for an overnight incubation at 4° C. The next day, sections were washed three times for 5 minutes in TBST, three times for 2 hours in TBST, three times for 10 minutes in AP buffer (NTMT; 0.1 M Tris-HCl pH 9.5, 0.1 M NaCl, 50 mM MgCl_2_, 1% Tween-20 in dH_2_O), and then incubated in the dark in NBT (nitro blue tetrazolium)/BCIP (5-bromo-4-chloro-3-indolyl-phosphate) staining solution (20 μL NBT/BCIP stock solution (Roche) per mL of NTMT) at RT. Once color development was complete, the NBT/BCIP solution was replaced with PBS to stop the staining reaction. Finally, sections were mounted on Superfrost Plus slides (Fisher Scientific) in Aqua-Poly/Mount (Polysciences).

Images were acquired with a camera-equipped Olympus IX51 inverted microscope using Cell^A (Olympus), and further processed with Adobe Photoshop CS6 or ImageJ (U.S. National Institutes of Health).

### Immunohistochemistry

Serial free-floating brain sections were obtained by slicing brains embedded in 4% w/v agarose/PBS with a vibratome (VT1000s, Leica) at 40 μm (E14.5 brains) or 60 μm (E13.5 brains) of thickness, and were collected in PBS. Tissue was permeabilized by washing three times for 10 (40 μm sections) or 15 (60 μm sections) minutes per wash in PBS-Triton X-100 0.2% at RT. Sections were then incubated in blocking buffer (10% solution of normal serum derived from the species in which secondary antibodies were raised in PBS) for at least 1 hour at RT, or overnight at 4° C.

Following blocking, the tissue was incubated with primary antibodies diluted in PBS-NaN_3_ 0.1% for 24–48 hours at 4° C. Primary antibodies used were mouse anti-neurofilament 2H3 (1:250; Developmental Studies Hybridoma Bank), rabbit anti-Islet1 (1:500; Abcam), mouse anti-Islet1 (1:50; Developmental Studies Hybridoma Bank), goat anti-mouse Sema6A (1:100; R&D), Armenian hamster anti-PlexinA4 (Mab-A4F5, 1:500; see Suto et al. [44]), Armenian hamster anti-PlexinA2 (Mab-A2D3, 1:50; see Suto et al. [41]). The monoclonal anti-NF 165 kD (2H3) and anti-Islet1/2 homeobox (39.4D5) antibodies, obtained from the Developmental Studies Hybridoma Bank (University of Iowa), were respectively developed by T.M. Jessell and J. Dodd, and by T.M. Jessell and S. Brenner-Morton.

Sections were afterwards washed three (40 μm sections) to five (60 μm sections) times for 10’ each in PBS at RT, and incubated for 2 hours at RT, or overnight at 4° C, with species-specific secondary antibodies conjugated with cyanine dyes (Jackson Immunoresearch) or Alexa Fluor^®^ dyes (Invitrogen) in a 1:500 dilution in PBS. They were next washed three to five times in PBS for 10 minutes each, and post-fixed in 1% PFA/PBS for at least 1 hour at RT, or overnight at 4°C. After the post-fixation step, sections were counterstained with 4’,6-Diamidino-2-Phenylindole (DAPI) (Invitrogen), and mounted on Superfrost Plus slides in Aqua-Poly/Mount. Slides were examined under an epifluorescence microscope (Axioplan2, Zeiss) connected to a digital CCD camera (DP70, Olympus) or a laser confocal microscope (LSM 700, Zeiss), and pictures acquired using analySISB (Olympus) or ZEN 2009 (Zeiss). Images were further processed and analyzed with Adobe Photoshop CS6 or ImageJ.

### Neuroanatomical tracing experiments

To back-label thalamic neurons projecting abnormally in the ventral telencephalon of *Plxna2;Plxna4* double mutant mice, small crystals of 1,1’-dioctadecyl-3,3,3’,3’-tetramethylindocarbocyanine perchlorate (DiI) (Molecular Probes) were inserted with a tungsten dissecting probe (World Precision Instruments) into superficial vTel layers of P0–P2 brain hemispheres. Back-labeling of thalamic neurons from either the primary visual (occipital) or primary somatosensory (parietal) cortex in *Plxna2;Plxna4* double mutant/wild-type mice was performed by respectively inserting small crystals of DiI and 4-(4-(dihexadecylamino)styryl)-N-methylpyridinium iodide (DiA) in the superficial cortical layers of P0-P2 brain hemispheres. Following the insertion of dye crystals, brains were kept in 1% PFA/PBS at RT in the dark for 4-6 weeks to allow the complete diffusion of the tracer in the axonal tracts and cell populations of interest.

Similarly, retrograde tracing to label guidepost cells in the vTel of wild-type, *Sema6a* mutant and *Plxna2;Plxna4* double mutant mice was carried out by insertion of small DiI crystals in the dTh of E13.5 brains, hemisected with a microsurgical knife (MSP) in order to expose the thalamus. Brains were subsequently kept in 1% PFA/PBS at RT in the dark for two weeks.

At the end of their incubation period, brains were embedded into 4% w/v agarose/PBS, and sectioned with a vibratome at 60 μm of thickness. Sections were counterstained with DAPI, mounted in Aqua-Poly/Mount onto Superfrost Plus slides, and analyzed using an epifluorescence microscope (Zeiss) not more than one day after sectioning, to avoid artifacts due to local dye diffusion at the surface of the sections. For guidepost cells labeling, Z-stacks (4 μm Z-step) were acquired with a laser confocal microscope (LSM 700, Zeiss) using the ZEN 2009 (Zeiss) imaging software, and further processed with ImageJ (U.S. National Institutes of Health) to obtain maximum intensity Z-projections of the vTel area.

## Results

### *Plxna2;Plxna4* double mutant mice show defects in subpallial TCA guidance

In order to analyze the development of thalamocortical connections in *Plxna2* and *Plxna4* mutant mouse brains, immunohistochemistry for the 165 kDa neurofilament subunit (clone 2H3, Developmental Studies Hybridoma Bank), a pan-axonal marker, was first performed on *Plxna2^−/−^, Plxna4^+/+^, Plxna2^+/+^, Plxna4^−/−^, Plxna2^+/−^;Plxna4^+/−^, Plxna2^+/−^;Plxna4^−/−^, Plxna2^−/−^;Plxna4^+/−^,* and *Plxna2^−/−^;Plxna4^−/−^* early postnatal littermate brains (n ≥ 3 for all genotypes analyzed) (Figure 1). No defective TCA phenotype was observed in either single mutants for *Plxna2* and *Plxna4,* nor in *Plxna2;Plxna4* double heterozygous mice (Figure 1A-C). However, *Plxna2^-/-^;Plxna4^-/-^* mice were found to present a defect in TCA pathfinding at the vTel that is strikingly like that observed in *Sema6a* mutants (Figure 1D). In all *Plxna2^−/−^;Plxna4^−/−^* specimens analyzed (P0–P2, n = 5), caudally-projecting TCAs were found to misproject ventrally in the vTel rather than turning laterally to enter the internal capsule, which these axons completely avoided (Figure 2A,B).

**Figure 1.**
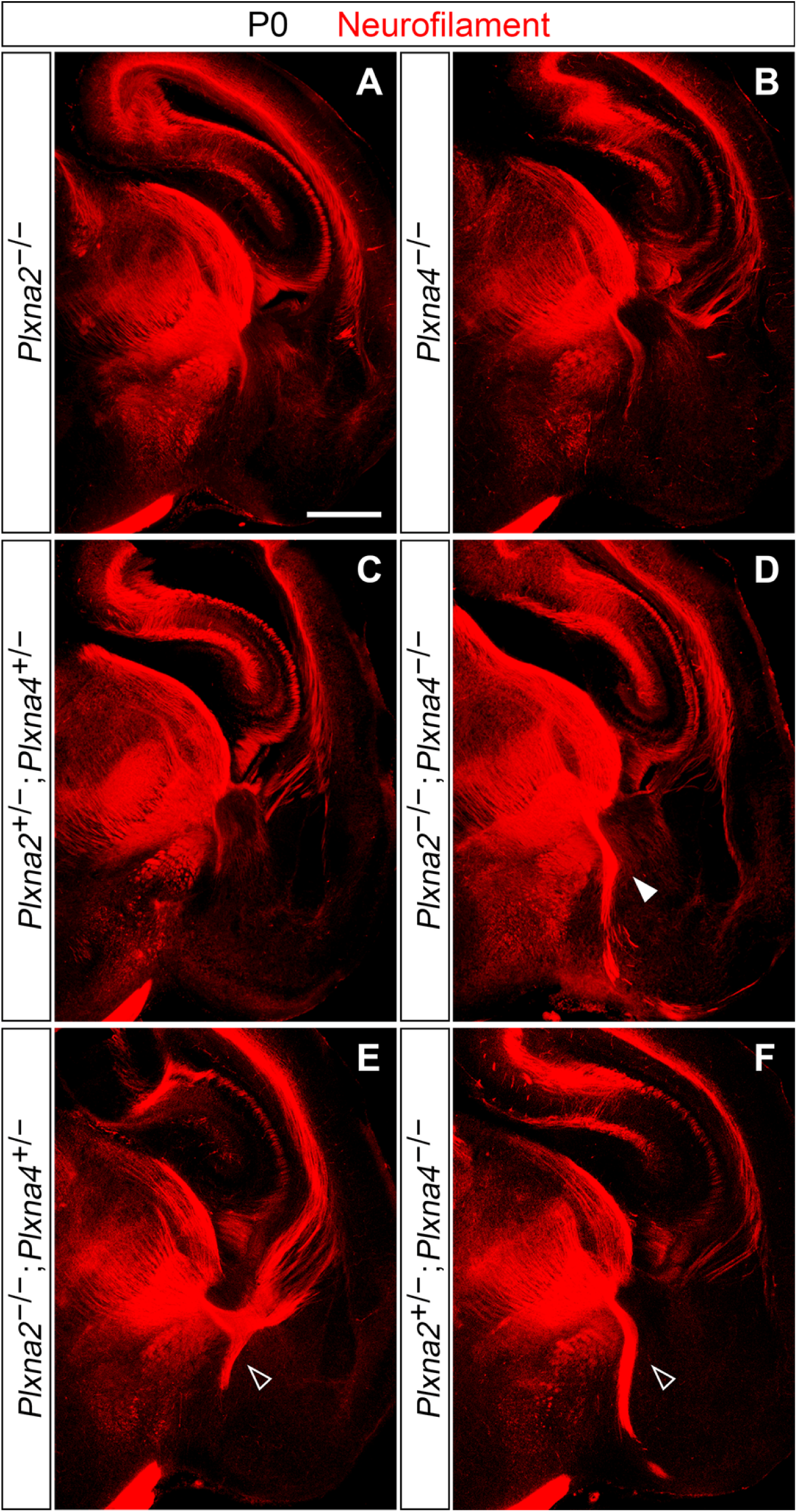
Immunohistochemical analysis of TCA development in *Plxna2;Plxna4* single and double mutant P0 brains. Immunostaining for neurofilament (red) confirms the presence of a Sema6a-mutant-like TCA defect in *Plxna2;Plxna4* double mutant postnatal brains (D; filled arrowhead), while no abnormal thalamic projections in the vTel are observed in either *Plxna2* or *Plxna4* single mutants (A, B). Additionally, misrouted TCAs are not present in *Plxna2;Plxna4* double heterozygous brains (C). Some TCA guidance defects, similar to those observed in double mutant brains but restricted to only a few thalamic fibers, also characterize *Plxna2^+/−^;Plxna4^−/−^* and *Plxna2^−/−^;Plxna4^+/−^* postnatal brains (E, F; empty arrowheads). Scale: 500 μm.

Detailed analysis of the trajectories followed by the misrouted TCAs also highlighted the presence, as in *Sema6a* mutants, of two discrete TCA bundles extending in distinct pathways in the vTel. At a more intermediate level of the rostro-caudal axis, axons were found to travel into the external capsule (Figure 2C-F), while more caudal TCAs, after their ventral turn, proceeded laterally within superficial vTel layers (Figure 2G-J). At P2, some of these axons were observed to extend along the pial edge of the vTel and reach the most superficial layers of the neocortex (Figure 2J), raising the possibility that, in later developmental stages, establishment of a normal thalamocortical connectivity through alternative routes might also occur in *Plxna2;Plxna4* double mutants.

**Figure 2.**
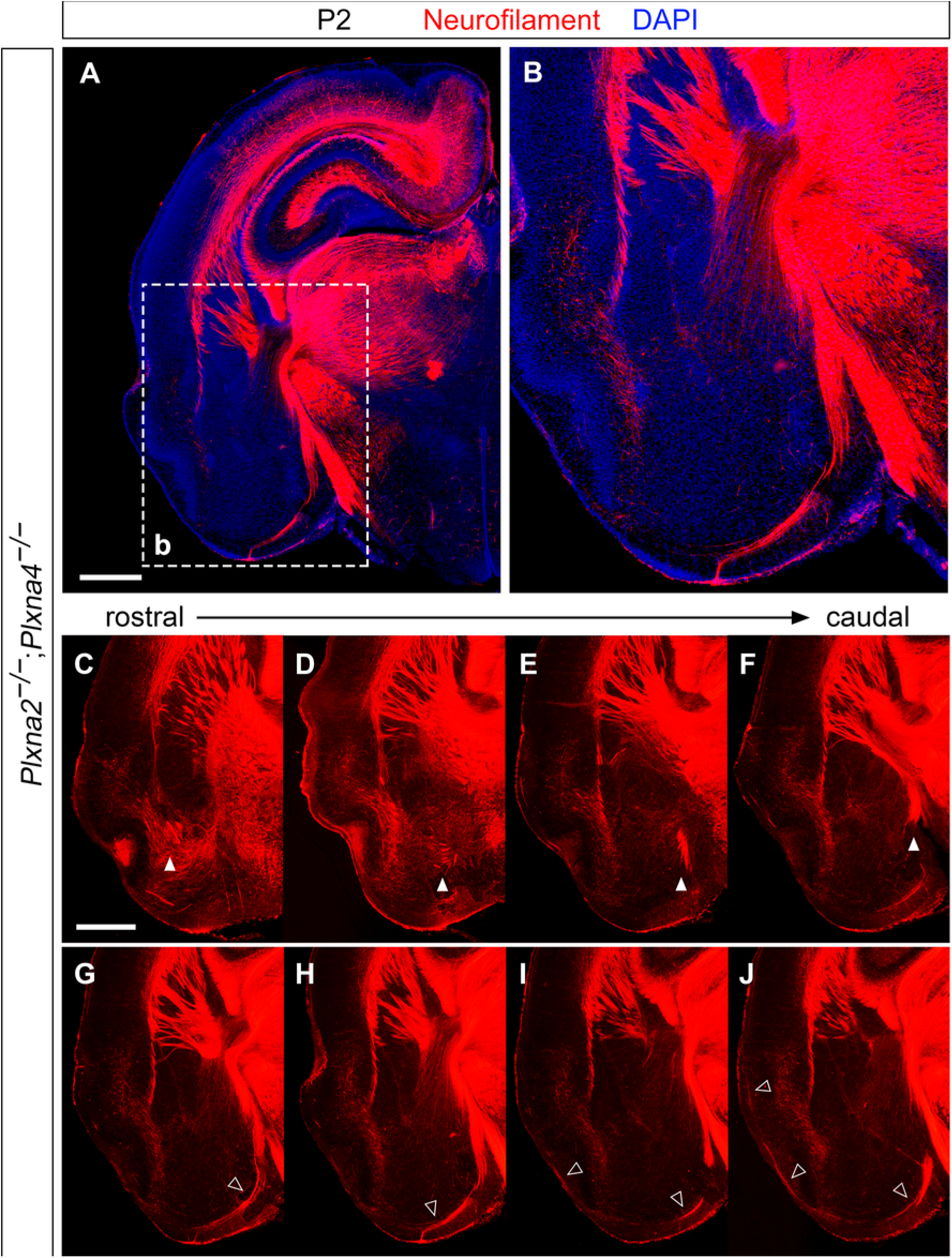
Immunohistochemical analysis of thalamocortical tract defects in a *Plxna2;Plxna4* double mutant P2 brain. Neurofilament immunostaining (red) reveals the presence of misrouted TCAs in the vTel of *Plxna2;Plxna4* double mutants (A, B). This TCA phenotype is markedly similar to that observed in *Sema6a* homozygous mutants; at more rostral levels (C–F), TCAs extend mainly along the external capsule (filled arrowheads), while at caudal levels (G–J) TCAs mostly follow a ventral route in the vTel, then proceed laterally across the telencephalon following a superficial pathway (empty arrowheads). Scale: A, 500 μm; B, 200 μm; C–J, 500 μm.

A compound effect of *Plxna2* and *Plxna4* loss-of-function mutations on thalamocortical connectivity was further demonstrated by the analysis of *Plxna2^+/−^;Plxna4^−/−^* and *Plxna2^−/−^;Plxna4^+/−^* mutant P0 brains: immunohistochemistry for neurofilament indeed revealed a few misprojecting caudal thalamic fibers in the vTel of these specimens (Figure 1E,F).

### Misrouted TCAs in *Plxna2;Plxna4* double mutants originate from dorsal lateral dTh nuclei

Retrograde neuroanatomical tracing methods were employed to identify the thalamic origin of the misrouted TCA bundles in the ventral subpallium of *Plxna2^−/−^;Plxna4^−/−^* P0 mouse brains (n = 4). As in *Sema6a* mutants, placement of DiI crystals at the vTel pial surface labeled thalamic projections which could be back-traced to the dLGN, as well as cell bodies within this nucleus (Figure 3A,B). In addition, *Plxna2;Plxna4* double mutants also showed back-labeling in axons and somas of a small, dorso-lateral portion of the VB (Figure 3B).

**Figure 3.**
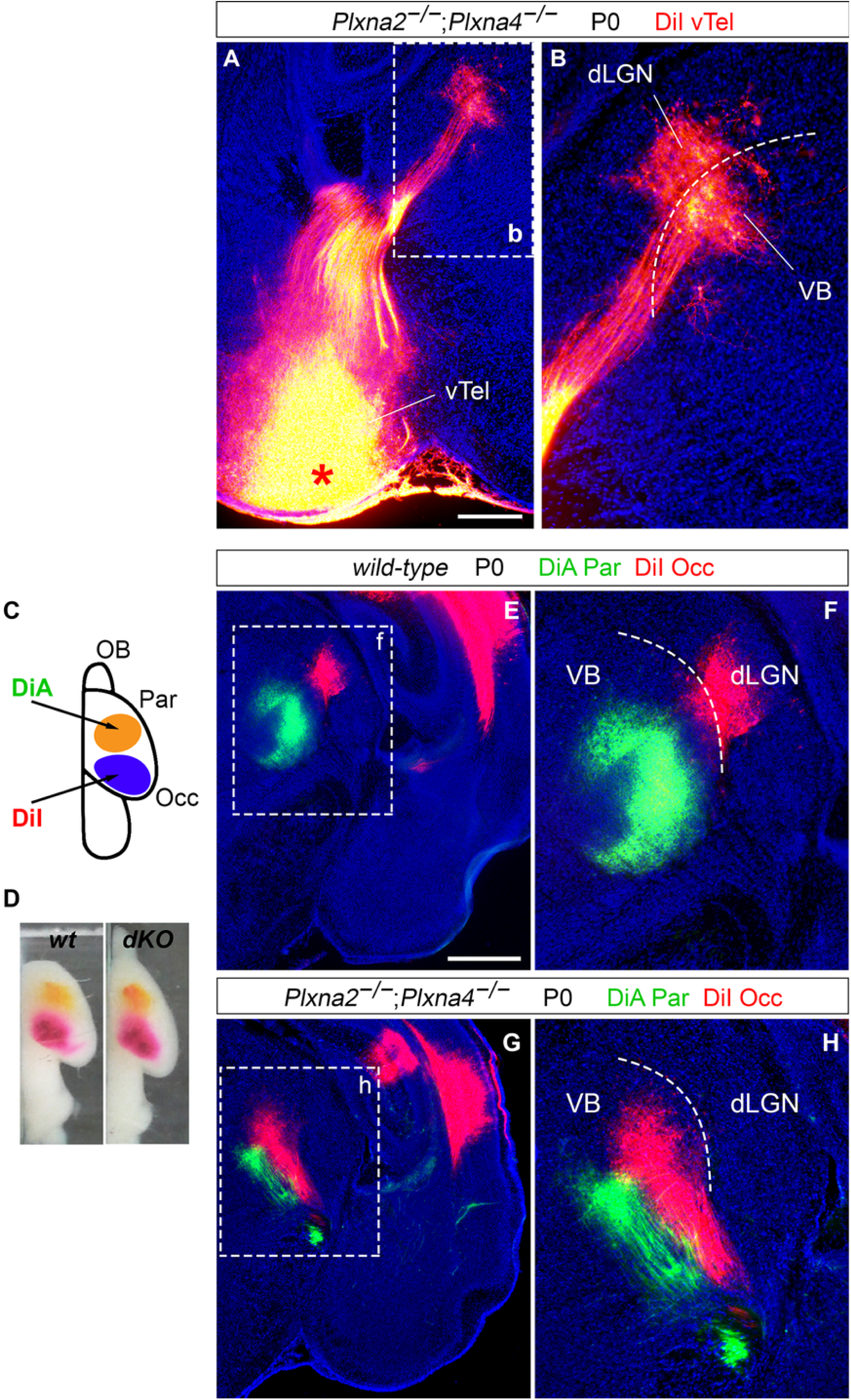
Neuroanatomical tracing experiments reveal a *Sema6a* mutant-like TCA phenotype in *Plxna2;Plxna4* double mutant brains. (A, B) Retrograde labeling with carbocyanine dyes from the vTel in the *Plxna2;Plxna4* double mutant P0 brain. (A) Insertion of DiI crystals in the vTel (asterisk) results in back-labeling of thalamic axons and cell somas located in the dLGN (B), a finding that coincides with data obtained from *Sema6a* mutant brains. In addition, dye-labeled neurons are also found in the VB, indicating the extension of guidance defects to a subset of thalamic axons normally directed to somatosensory cortical areas. (C–H) Back-labeling of thalamic neurons with two distinct carbocyanine dyes from the visual (occipital) cortex and the somatosensory (parietal) cortex in P0 wild-type (wt) and *Plxna2;Plxna4* double mutant (dKO) brains. (C) Schematic representation of the cortical sites of dye placement in P0 brain hemispheres (OB: olfactory bulb). DiA (green) and DiA (red) crystals are placed respectively on parietal (Par) and occipital (Occ) regions of the cortex. (E–H) Insertion of DiI crystals in visual cortical areas of *Plxna2^−/−^;Plxna4^−/−^* brains results in the back-labeling of some thalamic neurons of the dorso-lateral VB (H), rather than the dLGN (as instead observed in wild-type brains (F)), suggesting a miswiring of somatosensory TCAs to the visual cortex similar to that present in *Sema6a* mutants. Normal connectivity between ventro-medial VB neurons and the somatosensory cortex is preserved, as indicated by back-labeling of these cells by DiA. Scale: A, B: 250 μm; E–H: 500 μm.

### Somatosensory TCAs invade the visual cortex early postnatally in *Plxna2;Plxna4* double mutants

To explore the possibility of topographical alterations in the neocortex arising due to defects in subpallial dorso-lateral TCA guidance in *Plxna2^−/−^;Plxna4^−/−^* mice, retrograde double tracing experiments were performed from the visual (V1) and somatosensory (S1) cortex of *Plxna2^−/−^;Plxna4^−/−^* and wild-type P0 mouse brains (n = 4 per genotype) (Figure 3C-H). In *Plxna2^−/−^;Plxna4^−/−^* mice, back-labeling from V1 with DiI resulted in the identification of a subset of misprojecting VB neurons, indicating the invasion by somatosensory TCAs of this cortical region in the absence of dLGN projections (Figure 3G,H). Connectivity between some *Plxna2^−/−^;Plxna4^−/−^* somatosensory TCAs and their cognate cortical domains appeared to be preserved, as DiA crystals placed in the S1 led to the back-labeling of a ventro-medial VB cell population. Somas of the V1-invading thalamic neurons were observed adjacent ventro-medially to the VB neuronal subset that could be back-labeled from the vTel (Figure 3H), confirming the extension to somatosensory TCAs of the *Plxna2^−/−^;Plxna4^−/−^* miswiring phenotype characterized with vTel tracing experiments.

### Expression patterns of *Sema6a, Plxna2* and *Plxna4* during early TCA development

Overall, the phenotypical similarities observed between *Sema6a^−/−^* and *Plxna2^−/−^;Plxna4^−/−^* mutants provide evidence in support of a role of Sema6A–PlxnA2-PlxnA4 interactions in early dLGN axon guidance. In order to understand where and when these interactions may be required for proper subpallial TCA pathfinding, the spatiotemporal dynamics of *Sema6a, Plxna2* and *Plxna4* expression were first analyzed by *in situ* hybridization during TCA extension across the DTB and in the vTel.

Around embryonic day (E) 14.5, *Sema6a* was found to be expressed in all dorsal thalamic neurons, though more strongly in lateral regions, whereas *Plxna2* and *Plxna4* mRNAs were only detected in medial thalamic nuclei, notably excluding the region that will give rise to the dLGN (Figure 4A-D). At the same developmental stage, transcription of all three genes was observed in subpallial areas surrounding the IC, both in structures permissive for TCA growth, like the corridor, as well as non-permissive structures, such as the globus pallidus [22]. *Sema6a* mRNA was in addition detected at the vTel pial surface, the region invaded by misrouted TCAs in all of our mutant mice (Figure 4E-H) (n = 4 for each probe).

**Figure 4.**
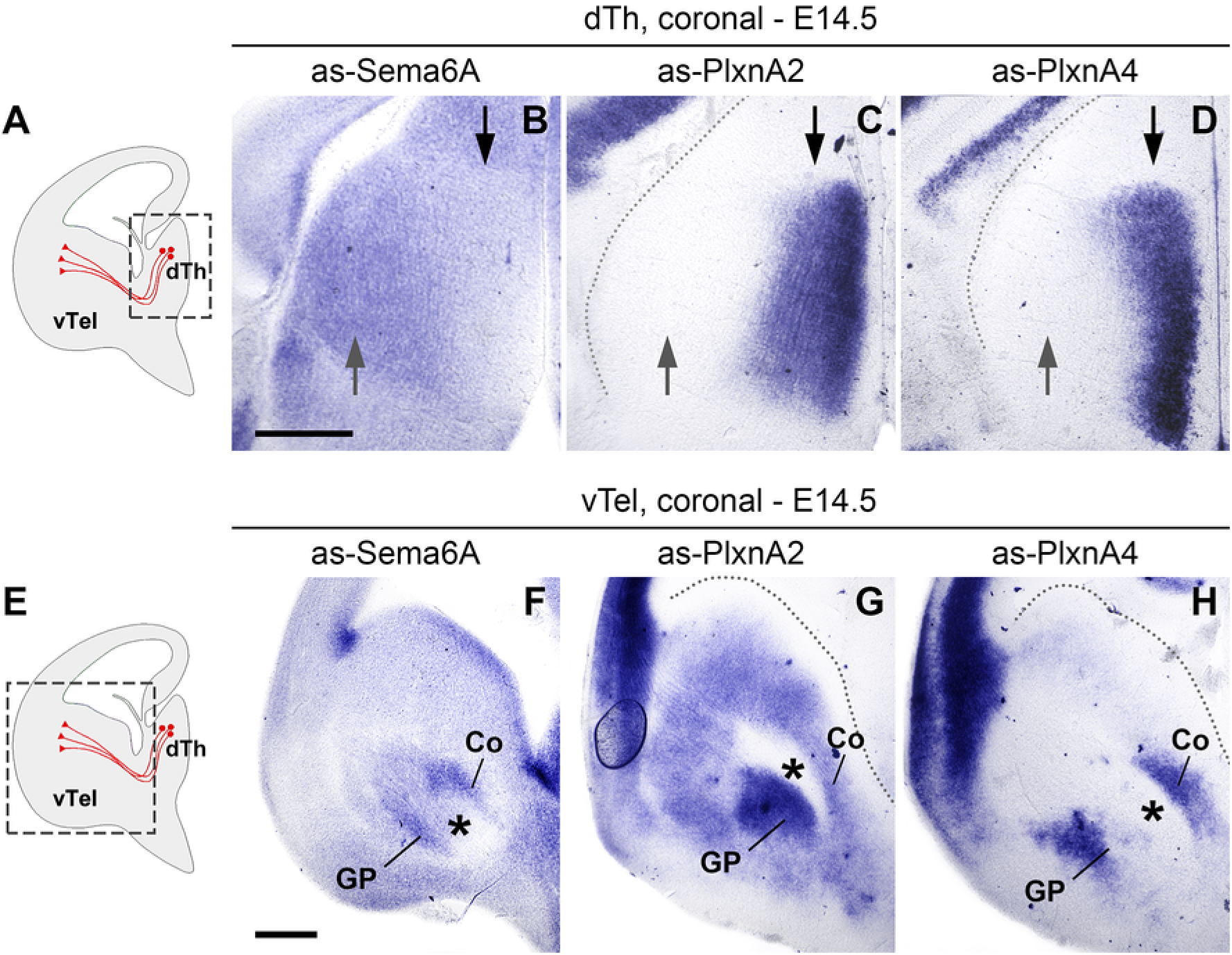
Expression of *Sema6a, Plxna2* and *Plxna4* mRNA in the E14.5 mouse forebrain. Levels of mRNA expression were detected by *in situ* hybridization with antisense (as) RNA probes. (A–D) mRNA expression in the dorsal thalamus (dTh) of E14.5 mouse brains. (A) Schematic representation of a coronal E14.5 section indicating the approximate region of interest (boxed area); TCAs are represented in red. (B-D) *Sema6a* is strongly expressed in both lateral (grey arrows) and medial (black arrows) dTh (B); in contrast, *Plxna2* and *Plxna4* are highly transcribed only in the medial dTh (C, D). (E–H) mRNA expression in the subpallium (vTel) of E14.5 mouse brains. (E) Schematic representation of a coronal E14.5 section indicating the approximate region of interest (boxed area); TCAs are represented in red. (F–H) *Sema6a* is highly expressed in areas surrounding the IC (asterisks), in the pial surface of the vTel, and the ventricular zone; it is also moderately transcribed in subventricular subpallial regions (F). *Plxna2* shows maximum expression in the globus pallidus, ventral to the IC, and strong expression levels in subventricular and mantle layers (G), while *Plxna4* is highly transcribed in two discrete bands located dorsal and ventral to the IC (H). Scale: A–D, 500 μm; E–H, 500 μm. Coronal section schemes adapted from López-Bendito et al. [22].

Overall, these findings suggest that Sema6A, PlxnA2 and PlxnA4 might be present on extending TCAs as they navigate subpallial territories, and furthermore be differentially expressed across axon subsets originating from different thalamic nuclei. Moreover, they indicate that these three guidance molecules are expressed from very early stages (at least as early as E12.5, data not shown) at the level of intermediate ‘decision points’ delineating TCA pathways in the vTel.

### Sema6A, PlxnA4 and PlxnA2 expression in thalamic neurons, TCAs, and the ventral forebrain in early TCA development

Based on *in situ* hybridization findings, double immunohistochemistry with neurofilament- and either Sema6A-, PlxnA2- or PlxnA4-specific antibodies on wild-type brains at E13.5 and E14.5 (n ≥ 3 per protein investigated) was next performed to analyze the spatiotemporal expression dynamics of these guidance molecules in distinct subsets of thalamocortical projections growing in the vTel.

At E13.5, Sema6A was found to be more highly expressed in dorso-lateral thalamic nuclei, while PlxnA4 and PlxnA2 expression appeared to be more restricted to medial thalamic nuclei; all proteins were observed along extending TCAs at moderate levels (Figure 5A-B’, Figure 5E-F’ and Figure S1, respectively). At E14.5, Sema6A was detected on all thalamic nuclei, but was found to be expressed only on the most caudally-located TCAs (which originate dorso-laterally in the thalamus) (Figure 5C-D’); PlxnA4 expression was observed only in medial thalamic nuclei and the rostrally-projecting TCAs originating from them, but some protein expression was surprisingly found on more caudally-located axons as well (Figure 5G-H’). However, at this caudal level the high degree of TCA fasciculation might not allow to completely distinguish projections directed to the somatosensory cortex from those directed to the visual cortex. Hence, PlxnA4 expression was also analyzed in *Sema6a* mutants at E15.5, a time-point at which misrouted thalamic fibers from the dLGN start to be clearly detectable as a bundle extending within the subpallial pial surface, completely detached from the IC. Immunostaining of tissue at caudal vTel levels revealed that, while PlxnA4 is present on TCAs elongating within the IC and presumably originating from the VB, this protein is not expressed by misprojecting dLGN axons (Figure 5I,J; n = 3). Taken comprehensively, these findings confirm *in situ* results at E14.5 showing differential expression in distinct thalamic nuclei.

**Figure 5.**
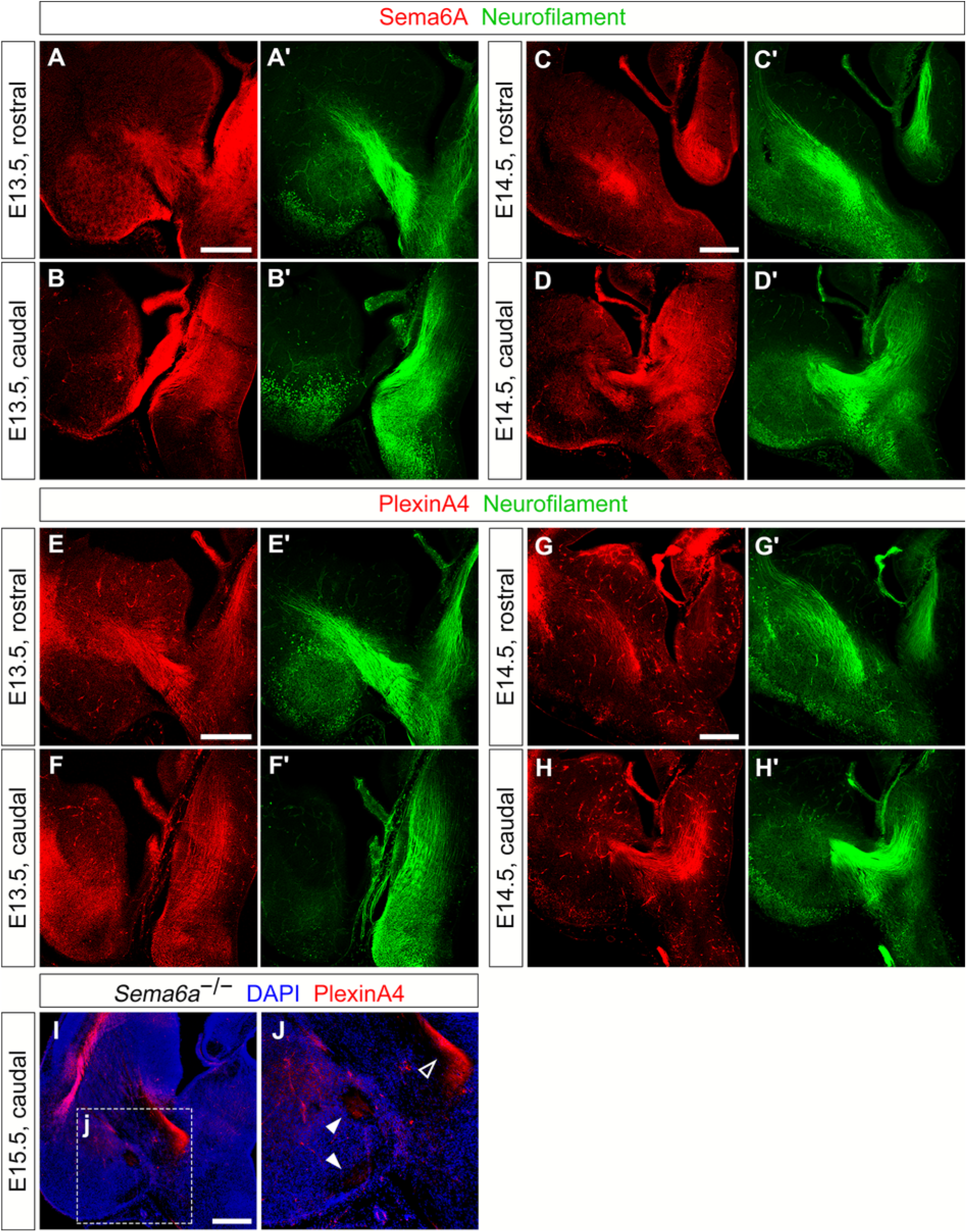
Expression of Sema6A and PlxnA4 on thalamic neurons and TCAs during axonal growth into the subpallium. (A–B’) Double immunohistochemistry for Sema6A (red) and neurofilament (green) on wild-type E13.5 coronal brain sections indicates expression of Sema6A in thalamic neurons, in particularly in dorso-lateral populations (B,B’), as well as on extending TCAs; the protein can also be found in vTel areas ventral to the IC (A,A’). (C–D’) Double immunohistochemistry for Sema6A (red) and neurofilament (green) on coronal sections of wild-type E14.5 brains shows expression of Sema6A extending to all thalamic nuclei (D,D’); the protein is furthermore expressed along TCAs positioned more caudally along the rostro-caudal axis (D,D’), while it is not found on TCAs projecting more rostrally (C,C’). (E–F’) Double immunohistochemistry for PlxnA4 (red) and neurofilament (green) in wild-type E13.5 coronal brain sections reveals that expression of PlxnA4 is mostly concentrated in medial thalamic neural populations, and is present on TCAs. Immunostaining can also be observed on some fibers in the IC contacting the axon bundle dorso-medially and ventro-laterally, and in mantle and pial surface areas of the caudal vTel (E,E’). (G–H’) Double immunohistochemistry for PlxnA4 (red) and neurofilament (green) on coronal sections of wild-type E14.5 brains demonstrates localization of PlxnA4 in medial thalamic neural populations (H,H’), as well as along TCAs projecting rostrally (G,G’). PlexinA4 seems to be further localized in some caudally-located TCAs (H,H’). (I–J) Immunohistochemistry for PlxnA4 (red) on coronal sections of *Sema6a^-/-^* E15.5 brains shows the presence of PlxnA4 on caudally-located TCAs extending within the IC (empty arrowhead), which presumably correspond to VB-originated axons. On the other hand, no immunostaining can be detected on misrouted projections corresponding to dLGN-originated fibers (filled arrowheads). Scale: A–B’, 300 μm; C–D’, 300 μm; E–F’, 300 μm; G–H’, 300 μm; I–J, 300 μm.

In addition, immunohistochemistry data highlighted areas of Sema6A, PlxnA2 and PlxnA4 expression in developing vTel domains surrounding the extending TCA bundle. For Sema6A, immunostaining could be detected in a restricted band dorsal to the IC, and in ventral domains extending to mantle and pial surface areas (particularly at intermediate / caudal levels) at E13.5; high expression appeared to additionally occur at the level of the lateral olfactory tract (Figure 5A). At E14.5, Sema6A could be moderately observed in areas possibly corresponding to corridor cell populations and in presumptive amygdalar territories; intense expression of the protein could be furthermore found at the level of the globus pallidus (Figure 5C,D).

For PlxnA4, immunostaining at E13.5 revealed the presence of this protein at intermediate vTel levels, in discrete domains surrounding dorso-medially and ventro-laterally the IC bundle, and a sparse expression in mantle / pial surface regions of the caudal vTel (Figure 5E,F). Staining could be observed also at E14.5 in a very small domain dorsal to the IC, but not in contact with the axons (Figure 5G).

For PlxnA2, immunostaining could be observed in a region likely overlapping the corridor at more intermediate levels of the E13.5 vTel; the protein was also found in medial-ventral areas of the caudal vTel (Figure S1A,B).

### Sema6A, PlxnA4, and PlxnA2 expression in vTel intermediate targets and guidepost neural populations

*In situ* hybridization and double immunohistochemistry experiments highlighted the expression of all our genes of interest in subpallial domains delineating TCA pathways in the vTel, and likely corresponding to structures crucial for proper guidance in these regions (e.g., the corridor). To better investigate spatiotemporal expression dynamics within intermediate TCA guidance targets, double immunohistochemistry for either Sema6A, PlxnA2 or PlxnA4 and Islet1, a marker for corridor cells and LGE-derived striatal neurons (Figure 6A), was performed on E13.5 (Figure 6B-H) and E14.5 (Additional file 1 - Figure S2, S3) wild-type brains (n ≥ 3 per experimental condition).

**Figure 6.**
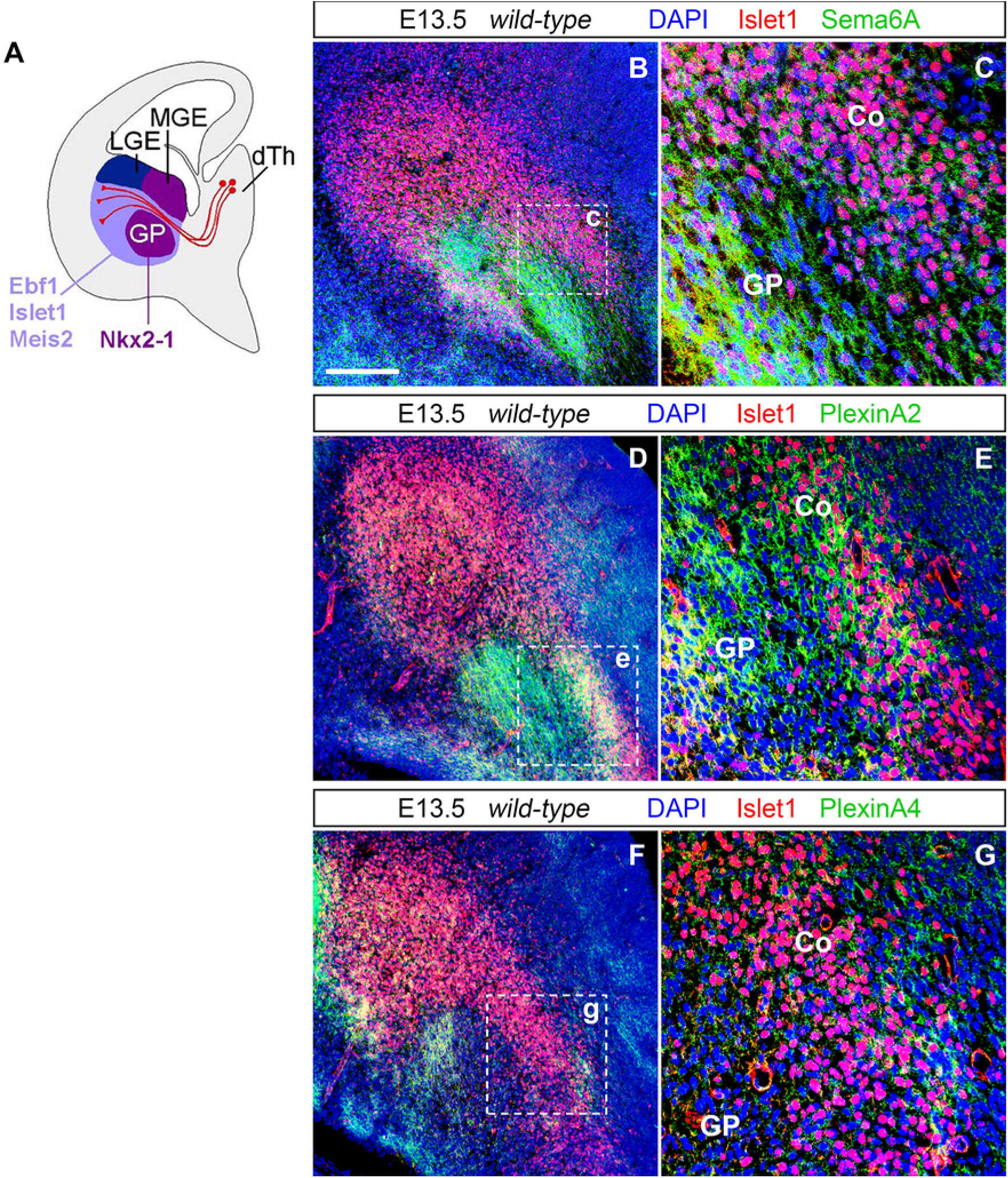
Expression of Sema6A, PlxnA2 and PlxnA4 in corridor cells and other subpallial structures at E13.5. (A) Diagram illustrating the spatial expression patterns of LGE- and MGE-derived neural population markers. The transcription factors Ebf1, Islet1, and Meis2 are detected in striatal and corridor regions of the vTel (light purple), both derivatives of the LGE, but not in the GP and the ventricular / subventricular zone of the MGE (dark purple). These territories in turn express a transcription factor, Nkx2-1, not present in LGE-derived territories. (Adapted from López-Bendito et al. [22].) (B, C) Double immunohistochemistry for the corridor cell marker Islet1 (red) and Sema6A (green) on coronal wild-type brain sections demonstrates the expression of Sema6A on corridor cells (Co) during the growth of TCAs into subpallial populations. Sema6A is also highly expressed in globus pallidus (GP) cells (C). (D–G) Double immunohistochemistry for PlxnA2 (green) and Islet1 (red) (D, E), or PlxnA4 (green) and Islet1 (red) (F, G) on coronal wild-type brain sections indicates a strong presence of PlxnA2 within the corridor and in the globus pallidus (E); PlxnA4 is also moderately present on most dorso-medial corridor domains (G), and in the lateral half of the globus pallidus area (F). Both molecules are additionally lightly expressed in the vTel subventricular zone and pial surface, in an area close to the IC, and in a discrete band at the ventral edge of the striatum (PlxnA4 is particularly present here) (D, F). Scale: 150 μm.

These experiments confirmed that Sema6A is consistently expressed during these developmental stages by corridor neurons and cells within the MGE-derived globus pallidus (Figure 6B-C). Furthermore, data indicated that Sema6A is present on most caudally-projecting TCAs, but also on other axonal tracts contained within the IC bundle: in particular, Sema6A appears to be expressed on nigrostriatal axons, that travel within the IC ventrally to and in close contact with TCAs (Figure S2).

Likewise, immunohistochemistry results for PlxnA2 confirmed the presence of this protein in corridor cells, and within the globus pallidus. PlxnA2 expression seems to extend, similarly to Sema6A, to multiple axonal projections extending within the IC more ventrally in respect to TCAs (Figure 6D-E). Concerning PlxnA4, immunohistochemistry data showed moderate expression of the protein in the corridor and in the globus pallidus at both E13.5 and E14.5 (Figure 6F-G, Additional file 1 - Figure S3). Compared to Sema6A and PlxnA2, PlxnA4 expression at E13.5 was observed in more limited domains overlapping with dorsal regions of the corridor, and lateral regions of the globus pallidus (Figure 6F). Interestingly, at E14.5 PlxnA4 was still found not only in the corridor, but also in discrete areas, particularly evident at caudal vTel positions, surrounding Islet-positive territories both dorso-medially and ventrolaterally (Additional file 1 - Figure S3).

Taken together, these results support a potential function of Sema6A, PlxnA2 and PlxnA4 not solely in the control of TCA guidance within subpallial territories, but also in corridor morphogenesis.

### Preserved corridor formation in *Sema6a* and *Plxna2;Plxna4* mutants

In order to investigate whether Sema6A, PlxnA2 and PlxnA4 might play a role in shaping the corridor domain, expression patterns for the corridor marker Islet1 in the vTel of wild-type, *Sema6a^−/−^, Plxna2^+/−^;Plxna4^+/−^* and *Plxna2^−/−^;Plxna4^−/−^* mice were examined at E13.5 and E14.5, at a time when LGE-derived neurons have terminated their migration while TCAs are extending into the IC, after crossing the DTB. Double immunohistochemistry for Islet1 and neurofilament on coronal brain sections revealed comparable patterns of the corridor cell marker’s expression in both striatal and IC regions between wild-type and mutant brains at E13.5 (Figure 7) and E14.5 (Additional file 1 - Figure S4) (n ≥ 3 for all genotypes in each experiment). At IC level, Islet1 was detected in a narrow band of cells lining a pathway for TCAs between the globus pallidus and the subventricular MGE zone, corresponding to the normal location of corridor neurons at these developmental stages. Additionally, immunostaining could be observed as normally expected in LGE-derived striatal territories, where TCA begin to rostro-caudally segregate in a fan-like shape. The local spatial distribution and density of corridor cells at E14.5 appeared to be fully preserved in all mutants (Additional file 1 - Figure S4, S5).

**Figure 7.**
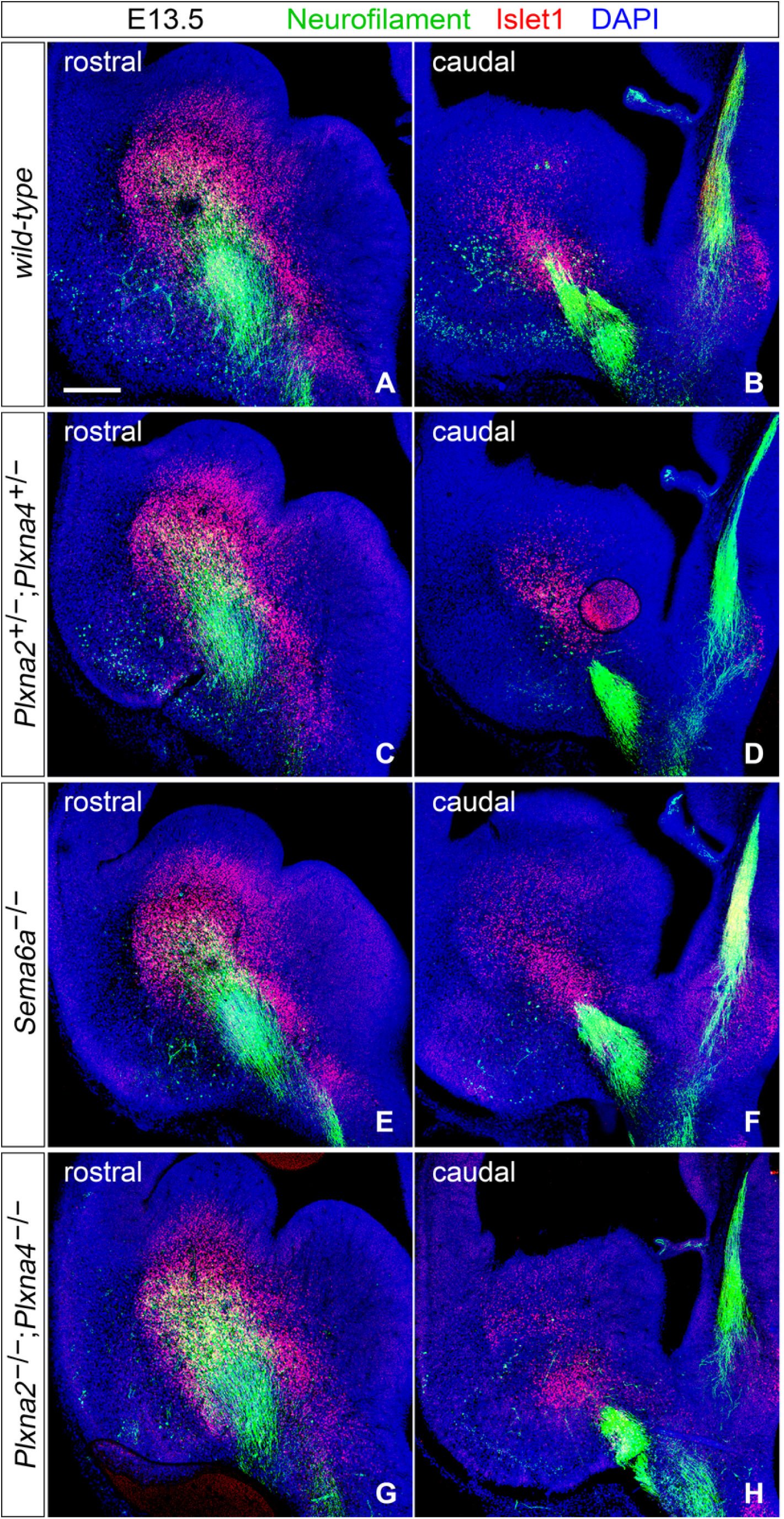
Normal overall expression of Islet1 in the vTel of *Sema6a* mutants and *Plxna2;Plxna4* double mutants at E13.5. Double immunohistochemistry for neurofilament (green) and Islet1 (red) on E13.5 coronal brain sections indicates the preserved organization, at this stage, of Islet1+ cell domains in the developing subpallium of *Sema6a^−/−^* (E, F), *Plxna2^+/−^;Plxna4^+/−^* (C, D), *Plxna2^−/−^;Plxna4^−/−^* (G, H) mouse brains, as compared to wild-type (A, B). Islet1+ neurons are present in a narrow band situated immediately dorsal to extending TCAs, between the vTel subventricular zone and the globus pallidus (characterized by the absence of Islet1 immunostaining), and throughout the striatum, where neurofilament-expressing thalamocortical fibers can be observed to segregate (A, C, E, G). Caudally, a slight reduction and disorganization of the most posteriorly-located subset of Islet1+ cells can be observed in the *Plxna2^−/−^;Plxna4^−/−^* mouse vTel (H). Scale: 200 μm.

Since dLGN axons are the first projections that cross the DTB during thalamocortical connectivity development, failure of pioneer caudally-directed axons to elongate with other thalamic fibers in the IC in *Sema6a-* and Plxna2;Plxna4-deficient mice could be due, for instance, to a delay in corridor cell migration. Thus, immunohistochemistry for Islet1 was performed on coronal brain sections of wild-type, *Sema6A^−/−^* and *Plxna2^−/−^;Plxna4^−/−^* E12.5 brains (when corridor formation is in its latest stages, and the first TCAs start crossing the DTB) to investigate the migration process of LGE-derived neurons to MGE-derived territories (n ≥ 3 for all genotypes). Comparison of Islet1 vTel expression pattern between wild type and mutant brain sections demonstrated the absence of any evident delay or corridor malformations: in all cases, Islet1+ cells could be distinguished in the mantle zone of the MGE-derived subpallial region (Additional file 1 - Figure S6).

Taken comprehensively, these findings suggest that loss of function of Sema6A, PlxnA2 and PlxnA4 does not impact on the overall development and spatiotemporal organization of the LGE-derived corridor and striatal populations in the mouse vTel.

### Misplacement of a subset of guidepost cells in *Sema6a* and *Plxna2;Plxna4* mutants

Guidance of TCAs across the subpallium has been suggested to rely not only on corridor cells, but also on IC-localized guidepost cells which form projections to the dTh just as TCAs start extending in the vTel (around E12.5). As there are no known molecular markers for these cells, they have been so far identified and studied by retrograde dye tracings experiments from the dTh [27]. Tracings with the carbocyanine dye DiI were therefore performed in *Plxna2^+/−^;Plxna4^+/−^* and *Plxna2^−/−^;Plxna4^−/−^* E13.5 brains from littermates to investigate whether loss of function of these guidance factors is associated with guidepost cell defects that might explain the TCA misrouting phenotype observed in *Plxna2;Plxna4* double mutants (Figure 8). In *Plxna2^+/−^;Plxna4^+/−^* brains, as expected, dye tracings from the dTh resulted in the back-labeling of cell bodies in the IC, in the proximity of anterograde-labeled TCAs and along the dorso-lateral pathway followed normally by TCAs during navigation into the subpallium (n = 7/7) (Figure 8D-G). In *Plxna2^−/−^;Plxna4^−/−^* brains, however, some back-labeled cell bodies were additionally found at a more superficial level of the vTel, in the presumptive amygdala; this group of cells was also observed more caudally in respect to the IC (n = 6/6) (Figure 8H-K).

**Figure 8.**
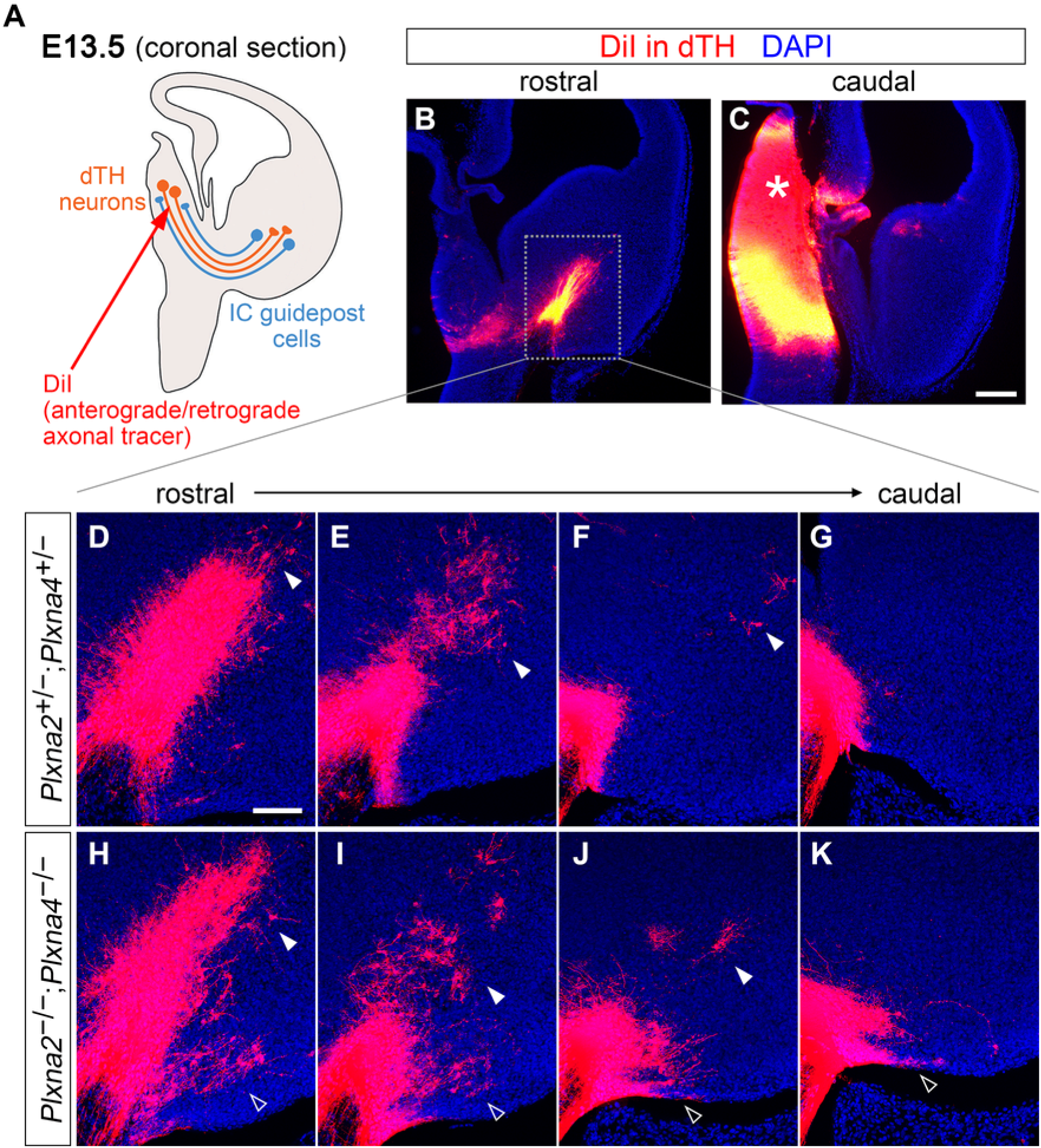
A subset of IC guidepost cells is misplaced in *Plxna2;Plxna4* double mutant E13.5 brains. (A) Schematic representation of the dye tracing experiments performed. DiI crystals were inserted into the dTh of E13.5 *Plxna2^+/−^;Plxna4^+/−^* and *Plxna2^−/−^;Plxna4^−/−^* mouse brains; from this position, the dye diffuses along TCAs in an anterograde fashion, and on guidepost cells’ projections reaching the dTh. (Adapted from Garel and López-Bendito [21].) (B, C) Coronal sections of E13.5 brains illustrating the labeled IC (B) and the exact location of dye placement in the dTh (asterisk in C). (D–K) DiI labels growing TCAs as well as guidepost cell bodies in the IC area, along the dorso-lateral path which TCAs will follow to proceed further into the subpallium (solid arrowheads), in both *Plxna2^+/−^;Plxna4^+/−^* (D-G) and *Plxna2^−/−^;Plxna4^−/−^* (H-K) brains. In *Plxna2^−/−^;Plxna4^−/−^* sections, however, back-labeling identifies a group of cells projecting to the dTh in an abnormal caudo-ventral position in the vTel (H-K, empty arrowheads). Scale: B, C: 250 μm; D-K: 100 μm.

Similar dye tracing experiments were next performed on *Sema6a^−/−^* E13.5 brains to examine the spatial distribution of IC guidepost cells in these mutants (late E13.5 wild-type brains (n = 10) were used as an extra control group). Like in *Plxna2^−/−^;Plxna4^−/−^* brains, along with somas normally localized in the IC region, some cell bodies were identified via back-labeling in posterior vTel surface areas of Sema6a-deficient specimens (these cells were not detected by retrograde tracing in wild-type brains) (n = 11/11) (Figure 9). In general, the positioning of these retrograde-labeled cells appeared to be less severely affected in *Sema6a^−/−^* brains; in some cases, only a very small number of cell bodies (n < 10) could be observed in aberrant sites throughout the vTel of these mutants (Figure 9I-L).

**Figure 9.**
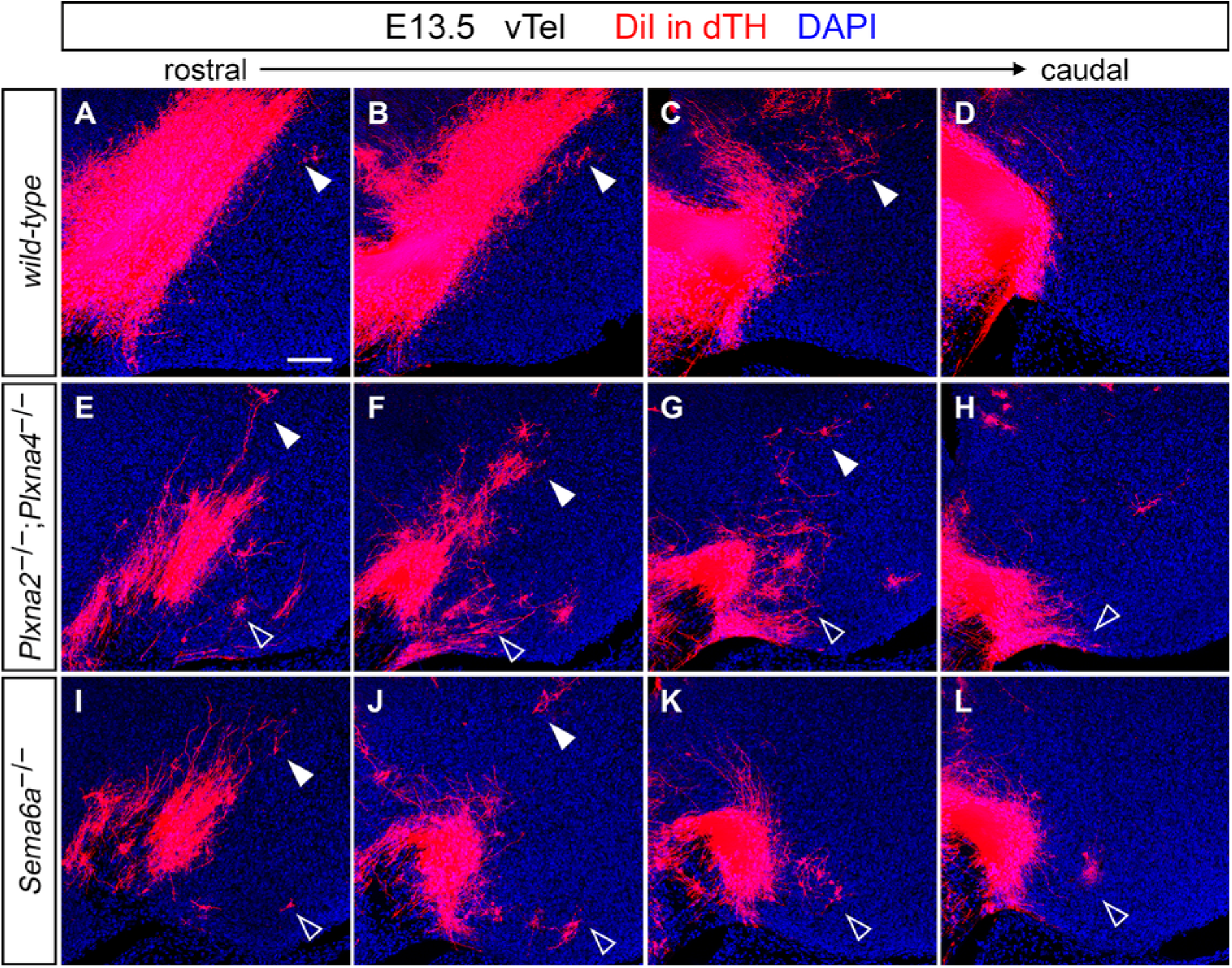
A subset of IC guidepost cells is misplaced in *Sema6a* mutant E13.5 brains. DiI retrograde tracing of cell populations within the vTel of wild-type, *Plxna2^−/−^;Plxna4^−/−^* and *Sema6a^−/−^* mouse brains reveals the presence in Sema6a-deficient mice of dTh-projecting neurons in caudal pial surface areas of the developing subpallium (I–L). No dye can be detected in this domain in late E13.5 wild-type brains (A–D), while back-labeled cells can be observed in the same region in *Plxna2^−/−^;Plxna4^−/−^* E13.5 mouse brains (E–H). (Images are taken at the same rostro-caudal positions in the mouse vTel as those in Figure 9D–K.) Scale: 100 μm.

Overall, these finding suggests that loss of Sema6A, PlxnA2 and PlxnA4 function in subpallial areas leads to the caudo-ventral misplacement of a subset of IC guidepost cells, which could explain the abnormal extension into the amygdala of more caudally-projecting TCAs.

## Discussion

### Sema6A, PlxnA2 and PlxnA4 act together in caudally-directed TCA navigation through the subpallium

Our investigation revealed that PlxnA2 and PlxnA4 act together in the guidance of the same thalamic fiber subset miswired in *Sema6a* null mutants, at the level of the same subpallial structures. As *Plxna2* and *Plxna4* single mutants do not show any defect in TCA guidance, the two proteins appear to have an at least partially redundant role in dLGN axon pathfinding. These results add to the body of evidence showing functional redundancy between Plexins, including PlxnA2/PlxnA4, in diverse neurodevelopmental contexts [37, 50, 51].

Furthermore, our findings highlighted an almost exact correspondence in several defects involving thalamocortical connections emerging as a result of either *Sema6A* or *Plxna2;Plxna4* ablation. Like in *Sema6a* null mutants, in *Plxna2^−/−^;Plxna4^−/−^* dLGN fibers turn ventrally once having crossed the DTB, instead of entering the IC, then proceed rostrally along the external capsule (eventually rejoining the IC axon bundle at the PSPB), or caudally within the pial surface of the vTel. Moreover, at P2 these latter misrouted axons can be observed to elongate dorsally in caudal pallial areas, at the level of the outermost neocortical layers. This finding might be indicative of an early postnatal recovery of visual thalamic nuclei–visual cortical areas connections via alternative axonal routes in superficial vTel and cortical territories, which is also characteristic of *Sema6a* mutants [34].

Another aspect of the *Plxna2^-/-^;Plxna4^-/-^* TCA pathfinding phenotype that mirrors that of *Sema6a^-/-^* mice is the invasion of visual cortical domains by somatosensory thalamic fibers originating from dorso-lateral VB regions. In Sema6a-deficient mouse brains, this caudal shift in TCA targeting is clearly observable around birth, but does not persist beyond P4, possibly due to somatosensory axons being out-competed by recovering visual projections in the innervation of the presumptive V1 [34]. Accordingly, the same shift in thalamocortical topography can be seen in P0 *Plxna2^−/−^;Plxna4^−/−^* mouse brains.

Interestingly, data here presented show that defects in TCA guidance in *Plxna2^−/−^;Plxna4^−/−^* mice extend to a small population of VB neurons located at the interface between visual and somatosensory thalamic nuclei. This observation suggests that additional cues might participate in PlxnA2/PlxnA4–dependent TCA guidance mechanisms, and indeed both Plexins have been shown to mediate signaling of other Semaphorin family members. The association of PlxnA2 with Semaphorin-6B and Semaphorin-5A plays an essential role during commissural and hippocampal axon guidance [44, 52, 53], and in the regulation of dentate gyrus granule cell synaptogenesis [54], respectively. As for PlxnA4, this guidance protein has been demonstrated to mediate axon-repulsive responses by binding to Semaphorin-6B [44], and to cooperate with other Semaphorin/Plexin cues in sensory and sympathetic axon pathfinding by interacting with Semaphorin-3A [44, 50]. PlxnA2 and PlxnA4 might therefore cooperate with additional Plexins in the subpallial guidance of axons from the VB.

Together with other findings generated by studies of mutant mouse lines presenting defects in TCA pathfinding within the basal forebrain, our results also support the notion that the switch from an external to an internal axonal path in the mammalian subpallium, due to changes in guidance cue patterning, is an evolutionarily recent trait connected with neocortical development in mammals [55]. Our mutant mouse lines specifically show that, in the event of a cytoarchitectural and/or functional disruption of structures delineating this internal path, thalamocortical connectivity can be eventually re-established following ancestral trajectories;however, this might occur at the expense of the proper functionality of the mature thalamocortical system [34, 56, 57].

### Complementary and overlapping *Sema6a, Plxna2* and *Plxna4* expression patterns in thalamic fibers and vTel guidepost cells during early TCA development

In this study, we expanded upon previous knowledge specific to late stages of TCA subpallial development by examining expression profiles of all our genes of interest throughout critical steps of TCA subcortical navigation. Our results confirmed previous findings obtained by *in situ* hybridization studies of Sema6A, PlxnA2 and PlxnA4 mRNA expression [34, 58, 59].

At E13.5 and E14.5, Sema6A, PlxnA2 and PlxnA4 present subpopulation-specific expression profiles in thalamic neurons and their projections, as well as the vTel. In the dTh, Sema6A is expressed broadly, in a high-dorsal to low-medial gradient, while PlxnA2 and PlxnA4 are observed restrictedly in medial thalamic nuclei. Notably, this region excludes the developing dLGN. These proteins are also present on TCAs; at E14.5 Sema6A is present on caudally-located axons, but absent in rostrally-projecting fibers, while PlexinA4 was observed to principally localize on these latter projections. Consistent with the in situ hybridization patterns, PlxnA4 immunoreactivity was absent from misrouted dLGN axons in *Sema6a* mutants. These data indicate that the effects on dLGN axon projections seen in *Plxna2;Plxna4* double mutants must be cell non-autonomous.

During TCA elongation in the subpallium, Sema6A, PlxnA2 and PlxnA4 are also present in several regions of the vTel, including structures, such as the corridor and the globus pallidus, suggested to play a role in directing TCA guidance during the first steps of subpallial axonal growth [21, 22, 27, 60, 61]. Combining these observations with recent evidence indicating that PlxnA2/PlxnA4–Sema6A reverse signaling occurs in vitro [36] and regulates axon guidance and targeting during murine optic system development [37], it can be hypothesized that PlxnA2 and PlxnA4 may act as a guidance cue for dLGN axons in the vTel, at the level of the IC. In this scenario, PlxnA2/PlxnA4 could provide, for instance, a repellent signal constraining axonal growth within the IC and acting specifically on Sema6A-positive dLGN projections. At the same time, PlxnA2/PlxnA4–positive axons originating from other thalamic nuclei might be unresponsive to subpallial Sema6A due to *cis* PlxnA2/A4–Sema6A interactions in those thalamic axon populations [36, 38].

On the other hand, the expression of all three proteins in the developing basal ganglia suggests that Sema6A–PlxnA2-PlxnA4 interactions might intervene in the correct patterning of these important ventral forebrain regions. Since all proteins are present in partially overlapping patterns in several subpallial domains, both forward and reverse signaling mechanisms might be here involved, and could be modulated by *cis* as well as *trans* binding events.

### Sema6A, PlxnA2 and PlxnA4 cooperate in morphogenetic processes involving caudal guidepost cell populations

Our analysis of vTel development in *Plxna2^−/−^;Plxna4^−/−^* and *Sema6a^−/−^* mice focused on two specific cell populations suggested by several studies to play a role in subpallial TCA guidance, the corridor cells and the IC guidepost cells.

In case of the LGE-derived corridor neurons, we observed an overall preserved spatial and temporal organization of these cells within basal forebrain territories of both *Sema6a-* and *Plxna2;Plxna4-deficient* mouse brains. Specifically, findings demonstrated that corridor cell migration proceeds normally in all null mutant brains, indicating that a cooperative action of Sema6A, PlxnA2 and PlxnA4 is not required during these early patterning events of the vTel. Moreover, the corridor domain’s cytoarchitectural features were found to be, for the most part, comparable between wild-type and mutant mouse brains at later developmental stages, when TCAs are elongating into presumptive basal ganglia regions of the vTel.

Considering IC guidepost cells, evident abnormalities in the spatiotemporal organization of this neural population were observed in both *Sema6a* and Plxna2;Plxna4-deficient mouse brains. At E13.5, when TCAs have begun their extension into the subpallium, guidepost cells could be back-labeled via retrograde dye tracing from the dTh in normal, IC-proximal positions in both *Sema6a^−/−^and Plxna2^−/−^;Plxna4^−/−^* mutant brains. However, in the posterior part of the vTel, a small population of retrograde-labeled cell bodies was observed in a more ventro-caudal site with respect to the IC, which corresponds to a presumptive amygdala area invaded by misprojecting TCAs in the examined mutants. This clearly suggests that Sema6A, PlxnA2 and PlxnA4 together play a role in specifying the proper localization of a caudal subset of dTh-projecting guidepost cells.

In relation to a potential cell non-autonomous mechanism of Sema6A action in visual axon guidance, it is worth to examine findings from previous studies on *Ebf1*, *Inpp5e* and *Lhx2* null mutant mice, which all show disruption of Sema6A subpallial expression profiles in concomitance with TCA pathfinding errors. The loss of *Sema6a* mRNA expression domains in the developing vTel of *Ebf1-* and *Inpp5e-*deficient mice is correlated to a disorganization of cell populations in the developing basal ganglia (which in this cases include corridor cells), and the ectopic growth of caudally-projecting TCAs in the presumptive amygdala [58, 59]. On the other hand, in *Lhx2* knockout mutants, *Sema6a* is over-expressed in the caudal vTel, guidepost cells cannot be back-labeled from the dTh in the IC region, and TCAs fail to extend in the vTel [62]. Our results are thus in line with evidence supporting a direct function of Sema6A in the morphogenesis of the IC, and provide new mechanistic insights on its role in the positioning of IC guidepost cells.

The fact that no Islet1-positive cells were observed in the vTel pial surface of our mutants, where some IC guidepost cells are found to mislocalize, supports the idea that at least part of this latter population may be unrelated to Islet1-positive corridor neurons [60, 61, 63, 64]. Indeed, it has been recently shown that some Dlx5/6-positive, Islet1-negative cells participate in forming the axonal bridge that allows TCAs to cross the DTB [63].

Our data is also in accordance with evidence showing that abnormalities, such as cell loss or misplacement, at the level of the most caudal parts of the corridor domain or the globus pallidus are associated with severe defects in TCA subpallial pathfinding (involving, in some cases, almost all fibers) [57, 60, 61, 65, 66]. Furthermore, these findings support the scaffolding model of axon guidance that has been proposed for IC guidepost cells [27, 63]. This said, it must be noted that a somewhat similarly compromised IC guidepost cells localization in concomitance to partial TCA guidance miswiring in the ventral subpallium has been described, so far, only in mice lacking the transcription factor Emx2 [67]. However, IC guidepost cells have yet to be exhaustively investigated in several mutant lines presenting an abnormal growth of caudal TCAs towards the presumptive amygdala.

## Conclusions

Taken comprehensively, results from our investigation of corridor and IC guidepost cells suggest that Sema6A–PlxnA2-PlxnA4 interactions may participate, by acting on the formation of intermediate guidance structures, in indirect mechanisms of TCA axon guidance at the level of the subpallium. Our findings do not exclude, however, a potential additional role of Sema6A reverse signaling in directing axonal growth with respect to PlxnA2/PlxnA4-expressing intermediate subpallial targets. Moreover, we cannot rule out the possibility that proper growth of these axons, or their responsiveness to guidance cues along their trajectories, might additionally be influenced by axon-axon interactions between TCA subsets differentially expressing Sema6A, PlxnA2 and PlxnA4.

## List of abbreviations

DAPI: 4’,6-Diamidino-2-Phenylindole
DiA: 4-(4-(dihexadecylamino)styryl)-N-methylpyridinium iodide
DiI: 1,1’-dioctadecyl-3,3,3’,3’-tetramethylindocarbocyanine perchlorate
dLGN: Dorsal lateral geniculate nucleus
DTB: Diencephalic-telencephalic boundary
dTh: Dorsal thalamus
GP: Globus pallidus
IC: Internal capsule
LGE: Lateral ganglionic eminence
MGE: Medial ganglionic eminence
PlxnA2: Plexin-A2
PlxnA4: Plexin-A4
PSPB: Pallial–subpallial boundary
RT: Room temperature
S1: Primary somatosensory cortex
Sema6A: Semaphorin-6A
TCA: Thalamocortical axon
V1: Primary visual cortex
VB: Ventrobasal complex
vTel: Ventral telencephalon

## Declarations

### Ethics approval

All animal procedures were approved by the TCD Animal Research Ethics Committee, and performed in accordance with Irish regulations on the use of animals for scientific purposes (Statutory Instrument No. 566 of 2002 and No. 543 of 2012) and institutional guidelines.

### Authors’ contributions

MDM and KJM conceived and designed the study. MDM performed all immunohistological and neuroanatomical tracing experiments. GEL and MDM performed *in situ* hybridization experiments. MDM and KJM analyzed and interpreted all data, and wrote the manuscript. All authors read and approved the final manuscript.

## Acknowledgements

We would like to thank Hajime Fujisawa’s group at Nagoya University, Japan, for providing the Plexin-A2- and Plexin-A4-specific antibodies that were employed throughout this study. We are additionally grateful to Jackie Dolan, Olivia Bibollet-Bahena, Daniel Shanley, Ash Watson, and Trinity College Dublin’s Bioresources staff for their precious technical assistance and useful comments.

## Additional files

Additional file 1: Supplemental Material (.pdf, 6.96 Mbs).

## REFERENCES

1. Van Battum, E.Y., S. Brignani, and R.J. Pasterkamp, Axon guidance proteins in neurological disorders. The Lancet Neurology, 2015. 14(5): p. 532–546.

2. Amaral, D.G., C.M. Schumann, and C.W. Nordahl, Neuroanatomy of autism. Trends in Neurosciences, 2008. 31(3): p. 137–145.

3. Ameis, S.H. and M. Catani, Altered white matter connectivity as a neural substrate for social impairment in Autism Spectrum Disorder. Cortex, 2015. 62: p. 158–181.

4. Belmonte, M.K., et al., Autism and abnormal development of brain connectivity. The Journal of Neuroscience, 2004. 24(42): p. 9228–9231.

5. Geschwind, D.H. and P. Levitt, Autism spectrum disorders: developmental disconnection syndromes. Current Opinion in Neurobiology, 2007. 17(1): p. 103–111.

6. McFadden, K. and N.J. Minshew, Evidence for dysregulation of axonal growth and guidance in the etiology of ASD. Frontiers in Human Neuroscience, 2013. 7: p. 671.

7. Wass, S., Distortions and disconnections: disrupted brain connectivity in autism. Brain and Cognition, 2011. 75(1): p. 18–28.

8. Barch, D.M., Cerebellar-thalamic connectivity in schizophrenia. Schizophrenia Bulletin, 2014. 40(6): p. 1200–1203.

9. Canu, E., F. Agosta, and M. Filippi, A selective review of structural connectivity abnormalities of schizophrenic patients at different stages of the disease. Schizophrenia Research, 2015. 161(1): p. 19–28.

10. Fornito, A. and E.T. Bullmore, Reconciling abnormalities of brain network structure and function in schizophrenia. Current Opinion in Neurobiology, 2015. 30: p. 44–50.

11. Friston, K.J. and C.D. Frith, Schizophrenia: a disconnection syndrome. Clinical Neuroscience, 1995. 3(2): p. 89–97.

12. Innocenti, G.M., F. Ansermet, and J. Parnas, Schizophrenia, neurodevelopment and corpus callosum. Molecular Psychiatry, 2003. 8(3): p. 261–274.

13. Karlsgodt, K.H., et al., Developmental disruptions in neural connectivity in the pathophysiology of schizophrenia. Development and Psychopathology, 2008. 20: p. 1297–1327.

14. Pantelis, C., et al., Early and late neurodevelopmental disturbances in schizophrenia and their functional consequences. Australian and New Zealand Journal of Psychiatry, 2003. 37(4): p. 399–406.

15. Pantelis, C., et al., Structural brain imaging evidence for multiple pathological processes at different stages of brain development in schizophrenia. Schizophrenia Bulletin, 2005. 31 (3): p. 672–696.

16. Woodward, N.D., H. Karbasforoushan, and S. Heckers, Thalamocortical dysconnectivity in schizophrenia. American Journal of Psychiatry, 2012. 169(10): p. 1092–1099.

17. Cao, M., et al., Imaging functional and structural brain connectomics in attention-deficit/hyperactivity disorder. Molecular Neurobiology, 2014. 50(3): p. 1111–1123.

18. Konrad, K. and S.B. Eickhoff, Is the ADHD brain wired differently? A review on structural and functional connectivity in attention deficit hyperactivity disorder. Human Brain Mapping, 2010. 31(6): p. 904–916.

19. Qiu, M.-g., et al., Changes of brain structure and function in ADHD children. Brain Topography, 2011. 24(3-4): p. 243–252.

20. Braisted, J.E., R. Tuttle, and D.D.M. O’Leary, Thalamocortical axons are influenced by chemorepellent and chemoattractant activities localized to decision points along their path. Developmental Biology, 1999. 208(2): p. 430–440.

21. Garel, S. and G. López-Bendito, Inputs from the thalamocortical system on axon pathfinding mechanisms. Current Opinion in Neurobiology, 2014. 27: p. 143–150.

22. López-Bendito, G., et al., Tangential neuronal migration controls axon guidance: a role for Neuregulin-1 in thalamocortical axon navigation. Cell, 2006. 125(1): p. 127–142.

23. Marín, O., et al., Guiding neuronal cell migrations. Cold Spring Harbor Perspectives in Biology, 2010. 2(2): p. a001834.

24. Raper, J. and C. Mason, Cellular strategies of axonal pathfinding. Cold Spring Harbor Perspectives in Biology, 2010. 2(9): p. a001933.

25. Squarzoni, P., M.S. Thion, and S. Garel, Neuronal and microglial regulators of cortical wiring: usual and novel guideposts. Frontiers in Neuroscience, 2015. 9: p. 248.

26. López-Bendito, G. and Z. Molnár, Thalamocortical development: how are we going to get there? Nature Reviews Neuroscience, 2003. 4(4): p. 276–289.

27. Molnár, Z., et al., Mechanisms controlling the guidance of thalamocortical axons through the embryonic forebrain. European Journal of Neuroscience, 2012. 35(10): p. 1573–1585.

28. Price, D.J., et al., The development of cortical connections. European Journal of Neuroscience, 2006. 23(4): p. 910–920.

29. Auladell, C., et al., The early development of thalamocortical and corticothalamic projections in the mouse. Anatomy and Embryology, 2000. 201(3): p. 169–179.

30. Garel, S. and J.L.R. Rubenstein, Intermediate targets in formation of topographic projections: inputs from the thalamocortical system. Trends in Neurosciences, 2004. 27(9): p. 533–539.

31. Leyva-Díaz, E. and G. López-Bendito, In and out from the cortex: development of major forebrain connections. Neuroscience, 2013. 254: p. 26–44.

32. Vanderhaeghen, P. and F. Polleux, Developmental mechanisms patterning thalamocortical projections: intrinsic, extrinsic and in between. Trends in Neurosciences, 2004. 27(7): p. 384–391.

33. Leighton, P.A., et al., Defining brain wiring patterns and mechanisms through gene trapping in mice. Nature, 2001. 410(6825): p. 174–179.

34. Little, G.E., et al., Specificity and plasticity of thalamocortical connections in Sema6A mutant mice. PLOS Biology, 2009. 7(4): p. e1000098.

35. Rünker, A.E., et al., Mutation of Semaphorin-6A disrupts limbic and cortical connectivity and models neurodevelopmental psychopathology. PLOS ONE, 2011. 6(11): p. e26488.

36. Perez-Branguli, F., et al., Reverse signaling by Semaphorin-6A regulates cellular aggregation and neuronal morphology. PLOS ONE, 2016. 11(7): p. e0158686.

37. Sun, L.O., et al., Functional assembly of accessory optic system circuitry critical for compensatory eye movements. Neuron, 2015. 86(4): p. 971–984.

38. Haklai-Topper, L., et al., Cis interaction between Semaphorin6A and Plexin-A4 modulates the repulsive response to Sema6A. The EMBO Journal, 2010. 29(15): p. 2635–2645.

39. Jongbloets, B.C. and R.J. Pasterkamp, Semaphorin signalling during development. Development, 2014. 141(17): p. 3292–3297.

40. Sun, L.O., et al., On and off retinal circuit assembly by divergent molecular mechanisms. Science, 2013. 342(6158): p. 1241974.

41. Suto, F., et al., Interactions between plexin-A2, plexin-A4, and semaphorin 6A control lamina-restricted projection of hippocampal mossy fibers. Neuron, 2007. 53(4): p. 535–547.

42. Faulkner, R.L., et al., Dorsal turning of motor corticospinal axons at the pyramidal decussation requires plexin signaling. Neural Development, 2008. 3: p. 21.

43. Rünker, A.E., et al., Semaphorin-6A controls guidance of corticospinal tract axons at multiple choice points. Neural Development, 2008. 3(1): p. 1–19.

44. Suto, F., et al., Plexin-a4 mediates axon-repulsive activities of both secreted and transmembrane semaphorins and plays roles in nerve fiber guidance. The Journal of Neuroscience, 2005. 25(14): p. 3628–3637.

45. Matsuoka, R.L., et al., Transmembrane semaphorin signalling controls laminar stratification in the mammalian retina. Nature, 2011. 470(7333): p. 259–263.

46. Bron, R., et al., Boundary cap cells constrain spinal motor neuron somal migration at motor exit points by a semaphorin-plexin mechanism. Neural Development, 2007. 2: p. 21.

47. Renaud, J., et al., Plexin-A2 and its ligand, Sema6A, control nucleus-centrosome coupling in migrating granule cells. Nature Neuroscience, 2008. 11(4): p. 440–449.

48. Zhuang, B.Q., Y.R.S. Su, and S. Sockanathan, FARP1 promotes the dendritic growth of spinal motor neuron subtypes through transmembrane Semaphorin6A and PlexinA4 signaling. Neuron, 2009. 61(3): p. 359–372.

49. Mitchell, K.J., et al., Functional analysis of secreted and transmembrane proteins critical to mouse development. Nature Genetics, 2001. 28(3): p. 241–249.

50. Yaron, A., et al., Differential requirement for Plexin-A3 and -A4 in mediating responses of sensory and sympathetic neurons to distinct class 3 Semaphorins. Neuron, 2005. 45(4): p. 513–523.

51. Schwarz, Q., et al., Plexin A3 and plexin A4 convey semaphorin signals during facial nerve development. Developmental Biology, 2008. 324(1): p. 1–9.

52. Andermatt, I., et al., Semaphorin 6B acts as a receptor in post-crossing commissural axon guidance. Development, 2014. 141(19): p. 3709–3720.

53. Tawarayama, H., et al., Roles of semaphorin-6B and plexin-A2 in lamina-restricted projection of hippocampal mossy fibers. The Journal of Neuroscience, 2010. 30(20): p. 7049–7060.

54. Duan, Y., et al., Semaphorin 5A inhibits synaptogenesis in early postnatal- and adult-born hippocampal dentate granule cells. eLife, 2014. 3: p. e04390.

55. Bielle, F., et al., Slit2 activity in the migration of guidepost neurons shapes thalamic projections during development and evolution. Neuron, 2011. 69(6): p. 1085–1098.

56. Gezelius, H., et al., Genetic labeling of nuclei-specific thalamocortical neurons reveals putative sensory-modality specific genes. Cerebral Cortex, 2016.

57. Lokmane, L., et al., Sensory map transfer to the neocortex relies on pretarget ordering of thalamic axons. Current Biology, 2013. 23(9): p. 810–816.

58. Garel, S., et al., The early topography of thalamocortical projections is shifted in Ebf1 and Dlx1/2 mutant mice. Development, 2002. 129(24): p. 5621–5634.

59. Magnani, D., et al., The ciliogenic transcription factor Rfx3 is required for the formation of the thalamocortical tract by regulating the patterning of prethalamus and ventral telencephalon. Human Molecular Genetics, 2015. 24(9): p. 2578–2593.

60. Jia, Z., et al., Regulation of the protocadherin Celsr3 gene and its role in globus pallidus development and connectivity. Molecular and Cellular Biology, 2014. 34(20): p. 3895–3910.

61. Uemura, M., et al., OL-Protocadherin is essential for growth of striatal axons and thalamocortical projections. Nature Neuroscience, 2007. 10(9): p. 1151–1159.

62. Lakhina, V., et al., Early thalamocortical tract guidance and topographic sorting of thalamic projections requires LIM-homeodomain gene Lhx2. Developmental Biology, 2007. 306(2): p. 703–713.

63. Feng, J., et al., Celsr3 and Fzd3 organize a pioneer neuron scaffold to steer growing thalamocortical axons. Cerebral Cortex (New York, NY), 2016. 26(7): p. 3323–3334.

64. Tuttle, R., et al., Defects in thalamocortical axon pathfinding correlate with altered cell domains in Mash-1-deficient mice. Development, 1999. 126(9): p. 1903–1916.

65. Mandai, K., et al., LIG family receptor tyrosine kinase-associated proteins modulate growth factor signals during neural development. Neuron, 2009. 63(5): p. 614–627.

66. Morello, F., et al., Frizzled3 controls axonal polarity and intermediate target entry during striatal pathway development. The Journal of Neuroscience, 2015. 35(42): p. 14205–14219.

67. López-Bendito, G., et al., Role of Emx2 in the development of the reciprocal connectivity between cortex and thalamus. Journal of Comparative Neurology, 2002. 451(2): p. 153–169.

